# Genetic differences in host infectivity affect disease spread and survival in epidemics

**DOI:** 10.1101/483602

**Authors:** Osvaldo Anacleto, Santiago Cabaleiro, Beatriz Villanueva, María Saura, Ross D. Houston, John A. Woolliams, Andrea B. Doeschl-Wilson

## Abstract

Survival during an epidemic is partly determined by host genetics. While quantitative genetic studies typically consider survival as an indicator for disease resistance, mortality rates of populations undergoing an epidemic are also affected by tolerance and infectivity (i.e. the propensity of an infected individual to transmit disease). Few studies have demonstrated genetic variation in disease tolerance, and no study has demonstrated genetic variation in host infectivity, despite strong evidence for considerable phenotypic variation in this trait. Here we propose an experimental design and statistical models for estimating genetic diversity in all three host traits. Using an infection model in fish we provide, for the first time, direct evidence for genetic variation in host infectivity, in addition to variation in resistance and tolerance. We also demonstrate how genetic differences in these three traits contribute to survival. Our results imply that animals can evolve different disease response types affecting epidemic survival rates, with important implications for understanding and controlling epidemics.

## Introduction

Infectious disease presents an enormous threat to animal and human populations, with epidemic outbreaks often causing high mortality due to insufficient disease control strategies. Over the last decades genetic and genomic studies have produced compelling evidence of substantial genetic variability in host response to infectious agents, potentially affecting epidemic risks and survival^1–8^. Nevertheless, remarkably little is known about how the genetics of individuals affects survival and disease spread at a population level. Quantifying the host genetic contribution to epidemic risk and severity remains a long-standing challenge in infectious disease research^9–11^.

Quantitative genetic studies tend to consider disease resistance as the primary and often sole host target trait for genetic disease control. The definition of disease resistance varies across studies depending on the disease in question and the knowledge of the host response mechanism exhibiting genetic variation^2^. However, due to large sample size requirements in quantitative genetic studies, disease resistance is defined by individual mortality, as it is easy to observe whether and when an individual exposed to infection dies. But survival is multifaceted, and may not only depend on the individual’s resistance to becoming diseased, but also on its *tolerance*, which in the context of survival may be defined as the propensity of an individual, once infected, to survive the infection^12,13^. Moreover, recently emerging evidence for the superspreader phenomenon, where a small proportion of highly infectious individuals are responsible for the majority of transmission, has raised awareness that individual variation in infectivity can also hugely influence population mortality rates^14–19^. Unsurprisingly, interest in the genetic regulation of infectivity, which can be defined as the host ability to infect an average susceptible individual upon unit contact, is increasing rapidly. Understanding the genetic regulation of infectivty is particularly pertinent if there are unfavourable genetic correlations between this trait and resistance or tolerance^17,20^.

Resistance, tolerance and infectivity may be regulated by different sets of genes with varying contributions, both in direction and in degree, to overall survival^21^. However, no study to date has simultaneously investigated these three traits. Whereas a plethora of quantitative genetic studies has demonstrated genetic variation in resistance^1,2,22–25^, relatively few studies have demonstrated genetic variation in disease tolerance^26–28^. No study to date has demonstrated genetic variation in host infectivity. Since genetic control strategies are environmentally friendly, non-intrusive and long-term preventive^29^, a more precise understanding of the genetic mechanisms underlying disease outbreaks and resulting mortality rates is urgently needed.

In this study, which considers infectious diseases with potentially fatal outcomes, we define disease resistance as an individual’s propensity for developing disease following contact with an infectious individual or material of average infectivity, and tolerance as the propensity of a diseased individual to survive. In line with these definitions, measurements of time of onset of disease and time from onset of disease to death in disease challenge experiments can disentangle resistance from tolerance. However, in the context of natural disease transmission, these phenotypes are no longer just affected by the genetics of the individuals in consideration. Instead they result from interactions between infected and non-infected individuals. Hence, time of onset of disease and time of death due to infection may no longer only depend on an individual’s own resistance and tolerance genes, but also on the infectivity genes of infected individuals in the same contact group. This complex genetic interaction calls for particular experimental designs and statistical models that can reliably disentangle and estimate genetic effects for all three epidemiological host traits.

Here we show how transmission experiments can be designed and analysed to identify genetic variation and disentangle genetic effects in resistance, tolerance and infectivity. We used infection by *Philasterides dicentrarchi* in turbot fish (*Scophthalmus maximus*) as a model system, for which genetic variation in mortality has been previously established^30^. *P. dicentrarchi* is a ciliate parasite which is the causative agent of scuticociliatosis, a common cause of mortality in turbot farming. Our resulting data not only allow quantitative genetic studies of infectious disease traits, but also enable evaluation of the relative contribution of resistance, tolerance and infectivity to survival. By analysing family differences in these traits, we demonstrate, for the first time, that there is genetic variance in infectivity, in addition to resistance and tolerance, and how genetic differences in host infectivity affect disease spread and survival.

## Results

### Designing disease transmission experiments to identify genetic variation in host resistance, infectivity and tolerance

Transmission experiments such as that depicted in Figure 1 provide data to identify genetic regulation of resistance, infectivity and tolerance to disease. Our experimental design meets the contrasting requirements for studying genetic regulation of each trait in isolation. In outbred populations, detection of genetic variation in host resistance and tolerance with respect to a particular pathogen can be deduced from family differences in response to infection, thus requiring related individuals from several families, ideally exposed to or infected with the same pathogen strain or dose. Genetic variation in these traits can then be detected through family differences in time to disease and time from disease to death, respectively. On the other hand, infectivity of a host impacts the disease status of its group mates rather than its own disease status. Because of this, and due to the difficulty in tracking who acquired infection from whom during an epidemic, host variation in this trait can be inferred by observing the speed at which the disease is naturally transmitted to susceptible individuals^31^. This can only be achieved by allocating genetically related infected individuals (shedders) across many contact groups containing susceptible individuals (recipients), such that recipient time to onset of disease can be observed (Figure 1). The groups should be closed such that transmission only occurs between shedders and recipients within the groups, therefore restricting the sources of infection and thus reducing uncertainty in infectivity estimates. Furthermore, in contrast to resistance and tolerance experiments, smaller groups are preferable to larger groups to minimise the confounding of the expression of infectivity from shedders and recipients^32,33^.

**Figure 1:**
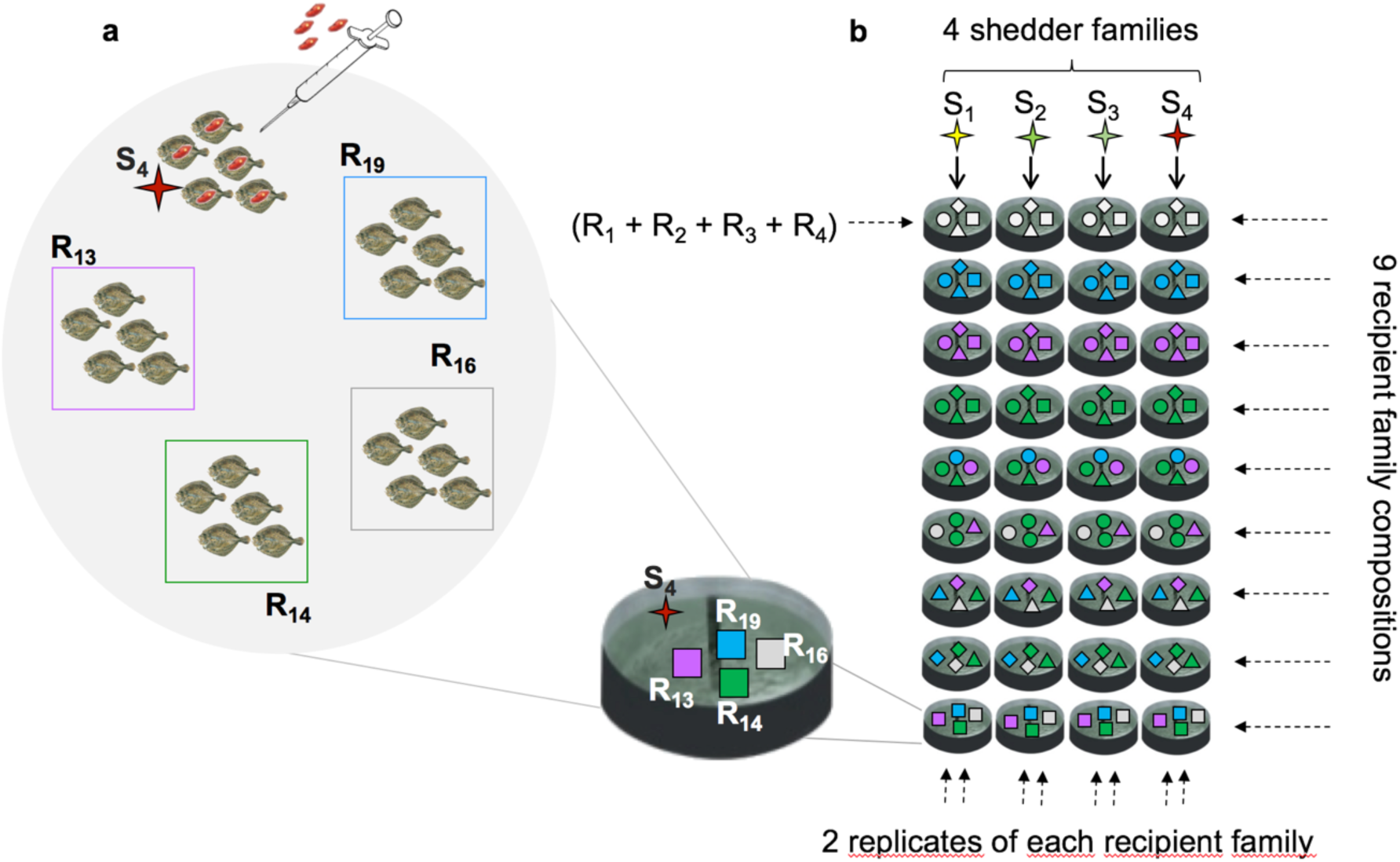
Transmission experimental design to detect genetic differences in host resistance, infectivity and tolerance. **a** Tank composition in our transmission experiment. Each of the 72 tanks comprised five artificially inoculated (shedder) fish from a single family and 20 susceptible (recipient) fish from four families. **b** Transmission experimental design in one of the trials. Each grey circle corresponds to one tank and a unique symbol-colour combination is assigned to each recipient (R_i_) or shedder (S_i_) family in 36 tanks in one of the two trials of our experiment. The 4 shedder families (S_1_ to S_4_) were distributed across nine tanks each. Recipient families were housed in tanks such that nine recipient family combinations were created, which in turn were housed with each of four shedder families. Each recipient family was allocated in two recipient family compositions.

Genetic differences in host infectivity based on shedder family information can be detected by analysing recipient time to onset of the disease. However, the risk of transmission and, consequently, time to onset of the disease depend not only on host infectivity but also on the resistance of recipient individuals^34^. Moreover, infected recipients may also become sources of transmission in closed groups. Therefore, to detect genetic differences in shedder infectivity, recipients exposed to different shedder families must have, on average, similar levels of resistance and infectivity. Then, a comparison of the speed of the transmission within recipients across shedders from different families allows detection of genetic variation in host infectivity. In particular, infection is expected to spread faster in groups of recipients exposed to individuals from a more infectious shedder family.

To obtain empirical data to study genetic variation in resistance, tolerance and infectivity, we performed a *Philasterides dicentrarchi* infection experiment in two large cohorts of turbot fish (total n = 1,800) from 44 full-sib families (see Figure 1). The visually conspicuous infection signs caused by this parasite enable non-invasive tracking of the epidemics in real-time^35–37^ (see Methods), which is usually difficult to observe in transmission trials. We distributed shedder fish artificially inoculated with the parasite across 72 isolated tanks, such that these fish transmitted the disease to previously non-infected recipient fish (see Methods).

Each tank comprised five shedder fish from the same family and 20 recipient fish from four different families, forming a recipient family composition (Figure 1). This group size and composition allowed to meet requirements for reliably detecting genetic differences in infectivity (which requires small groups with families distributed across groups^38,38^) as well as in recipient resistance and tolerance (which are best inferred from large groups comprising multiple offspring from many families). As environmental conditions were controlled (see Methods), the only sources of variation in tanks were recipient family composition and shedder family. Family composition in tanks was designed to detect genetic variance in host infectivity while having a suitable experimental setup to verify genetic differences in resistance and tolerance. To achieve this, we exposed each of four shedder fish families to the same nine recipient family compositions, in each of two trials of the experiment (see Methods). With this, infection was expected to spread consistently faster in recipient groups exposed to the most infectious shedder family, thus avoiding confounding between shedder family infectivity and genetic resistance and infectivity of recipient fish. Also, we included each recipient family in two recipient family compositions (Figure 1, see Methods). This way, each recipient family was exposed to all shedder families (to avoid confounding between recipient family resistance and shedder family infectivity) and a sufficiently large number of fish from different families, ensuring reliable detection of differences in resistance and tolerance between recipient families. Therefore, the resulting design allowed not only detection of genetic variation in infectivity through differences in disease onset profiles of recipient fish pooled by shedder families, but it also enabled detection of genetic variation in resistance and tolerance by comparing different profiles for onset of disease and infection-induced death, respectively, pooled by recipient family.

Disease and survival data from the experiment were collected by inspecting all fish twice a day over the duration of the experiment for visual signs of infection and for mortality. For each fish displaying visual infection signs, the date and time of first detection of signs was recorded. Similarly, for fish that had succumbed to the infection, the date and time of detected mortality was recorded. Whilst measurements of time to signs allow genetic analysis of resistance of recipient fish and infectivity of shedder fish, measurements of the time from signs to death provide information on genetic variation in tolerance.

### Analysis of time-to-event measurements associated with fish onset of disease and subsequent survival

It took longer for the recipients in the experiment to develop disease signs than to die following infection, with large variation in time to signs across trials (median time to signs was 43 and 110 days for trials 1 and 2 respectively, see also Figure 2). This large variation between trial 1 and 2 might be caused by lower virulence of the pathogen strain in trial 2, as a later onset of visual signs in recipient fish were found in that trial. Correspondingly, trials 1 and 2 lasted respectively 104 and 160 days (see Methods). There was however similar within-trial variation in time to signs and time from signs to death, suggesting that the same traits were measured despite the different pathogen virulence in both trials. (Figure 2). Fish from both trials died quickly and with low variation following onset of signs (Figure 2). Median times from signs to death were 8 and 7 days for recipient fish from trials 1 and 2 respectively. Also, the large recipient variation in time to signs that we could not observe in time from signs to death can be explained by factors associated with tank composition as described below.

**Figure 2:**
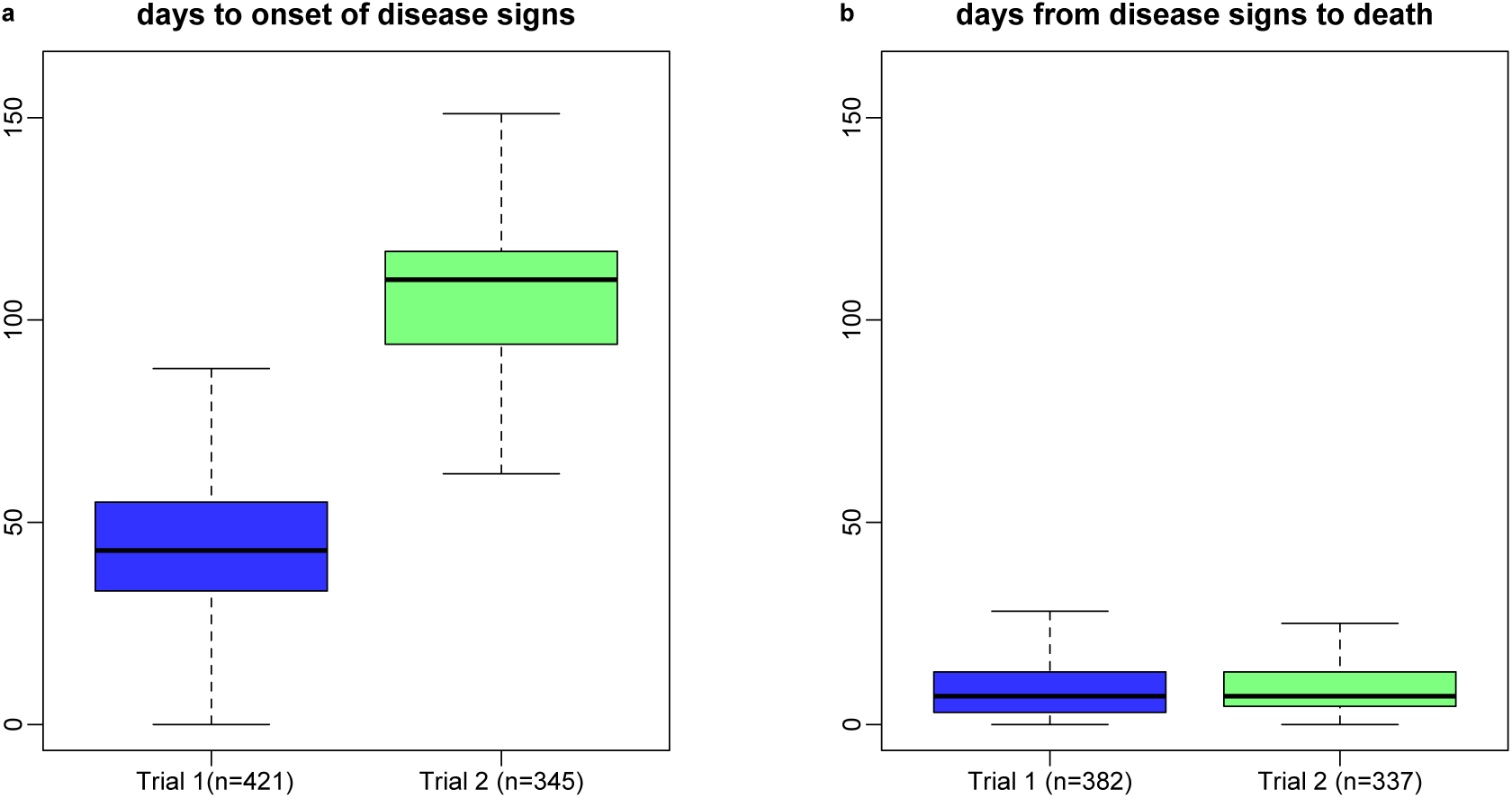
Turbot might have a short lifespan following infection by Philasterides dicentrarchi. **a** Box plots of time to disease signs for recipient fish that displayed these signs in the two trials of our experiment. **b** Box plots of time from signs to death for recipient fish considered in (a) that died before the end of the experiment.

Long recipient survival post exposure was strongly associated with long time to signs (Kendall correlation coefficients 0.70 and 0.74 for trials 1 and 2 respectively, P< 0.001, see Figure 3), but weakly correlated with time from signs to death (Kendall correlation coefficients are 0.20 (P < 0.001) and −0.01 (P = 0.50) for trials 1 and 2, respectively). We also found a weak positive relationship between recipient time to signs and time from these signs to death (Kendall correlation coefficients −0.19 (P = 0.003), and −0.34 (P < 0.001) for trials 1 and 2, respectively, see also Figure 3). These results suggest that factors controlling the onset of disease have a stronger influence on survival than factors driving within host disease progression.

**Figure 3:**
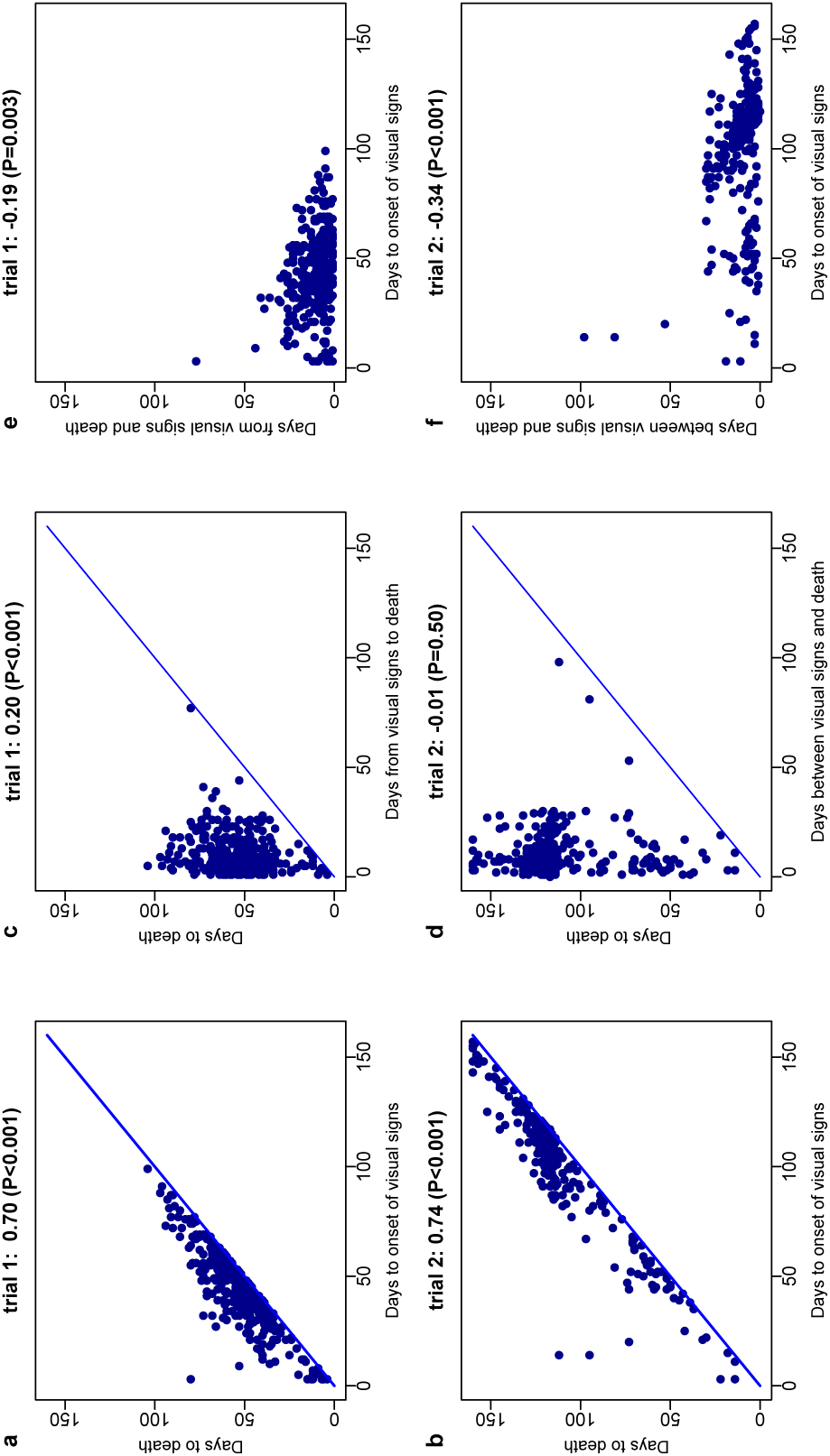
Turbot survival during disease outbreaks caused by *Philasterides dicentrarchi appear to be strongly influenced* by time to infection but weakly correlated with time from infection to death. Scatterplots and Kendall correlations between time-to-event measurements collected for recipient fish that showed disease signs and died during trials 1 (n=382) and 2 (n=337) of the transmission experiment. **a** and **b** time to death *vs* time to signs (distance between dots and diagonal line represents time from signs to death). **c** and **d** time to death *vs* time from signs to death (distance between dots and diagonal line represents time to signs). **e** and **f** time from signs to death *vs* time to signs. P-values were calculated using Kendall’s tau statistic (two-tailed).

### Fish family variation in disease resistance and tolerance

We found significant differences across the 36 recipient fish families in time to signs (log-rank test: P < 0.001 for both trials) but small recipient family variation in time from signs to death (log-rank test: P = 0.053 and P = 0.084 for trial 1 and 2, respectively, also see Figure 4). These results may indicate large genetic variation in resistance but small variation in tolerance to the disease. Large and low variation for, respectively, time to signs and time from signs to death were also found across tanks (Supplementary Figure 1).

**Figure 4.**
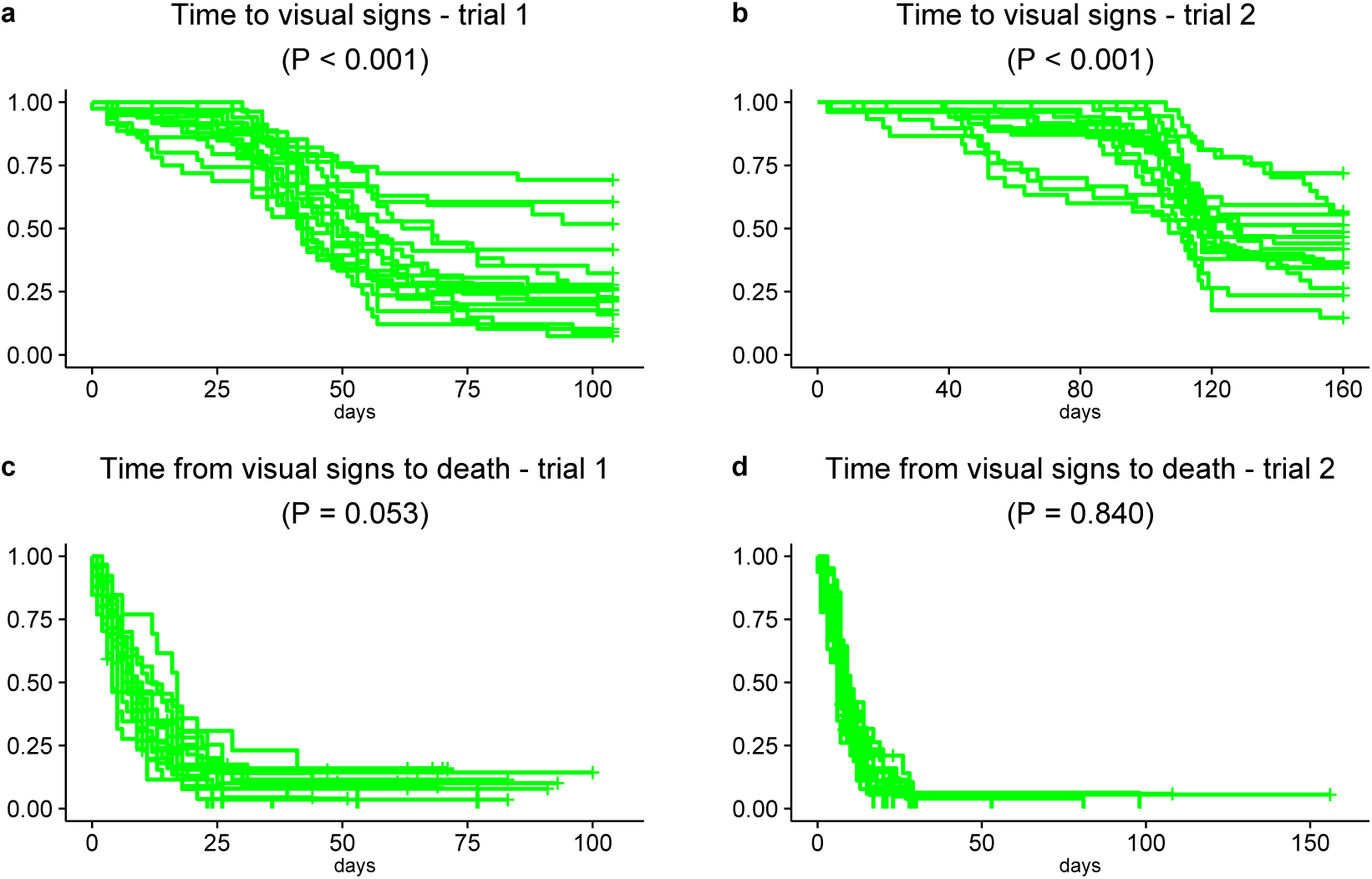
Family differences in onset of visual signs and survival post disease. Evolution of disease caused by *Philasterides dicentrarchi* (a-b) and survival post disease (c-d) in all families of recipient fish in trials 1 and 2 of the transmission experiment. The curves were obtained through family-based Kaplan-Meier plots for time to signs (a-b) and time from signs to death (c-d). P-values were calculated using the two tailed log-rank test for detecting family differences in Kaplan-Meier estimates. Number of fish per family for each of the graphs are presented in Supplementary tables 2 and 3.

### Effect of shedder family infectivity on onset of disease and subsequent survival

Shedder fish family had a strong effect on recipient time to signs but little influence on time from these signs to death (Figure 2 and Table 1). To quantify the effect of shedder family on recipient infection and survival, we fit generalized linear mixed models (GLMMs) to daily counts of recipient fish with visual signs and recipient fish that died due to infection (see Methods). When applied to discrete time-to-event data and assuming a baseline hazard for each day, these models can estimate hazard ratios between different shedder families while controlling for other factors of the transmission experiment^39,40^. Recipients exposed to the most infective shedder families (C and F for trials 1 and 2 respectively, see Figures 5a and 5b) got diseased at approximately twice the rate of recipients exposed to the least infective shedder families (B and G for trials 1 and 2 respectively, see Figures 5a and 5b; hazard ratio estimates: trial 1 posterior mean: 1.80, 95% CI: 1.37, 2.23; trial 2 posterior mean: 2.10, 95% CI: 1.60, 2.75). As a further illustration of the effects of genetic differences in shedder infectivity, we found that the relative hazard of a recipient showing disease signs when exposed to the most vs the least infective shedder family was greater than 1.5, with Bayesian posterior probabilities 0.85 and 0.98 for trials 1 and 2, respectively. In contrast, shedder family infectivity did not significantly affect recipient time of death following disease (hazard ratio estimates: trial 1 posterior mean: 1.27, 95% CI: 0.88, 1.84; trial 2 posterior mean: 0.98, 95% CI: 0.76, 1.21; Bayesian posterior probabilities that recipient hazard ratio of death following disease between most and least infective shedder fish family is at least 1.5: <0.005 for both trials).

**Table 1:** Effect of shedder fish family on time to signs and time from signs to death estimated with the Watanabe*-*Akaike information criterion (WAIC, see Methods) for generalised linear mixed models (GLMM) fitted to time-to-event measurements with and without shedder family as covariate. Inclusion of the shedder family as model covariate resulted in better predictive ability in GLMMs for time to onset of visual signs when compared to model predictive improvement due to inclusion of shedder family into the same models for time between infection signs and death.

| **Trial** | **response variable** | **GLMM with shedder family** | **GLMM without shedder family** |
| --- | --- | --- | --- |
| 1 | time to signs | 2419.33 | 2453.33 |
|  | time from signs to death | 1588.29 | 1590.4 |
| 2 | time to signs | 2365.69 | 2391.38 |
|  | time from signs to death | 1343.7 | 1357.54 |

**Table 2:**
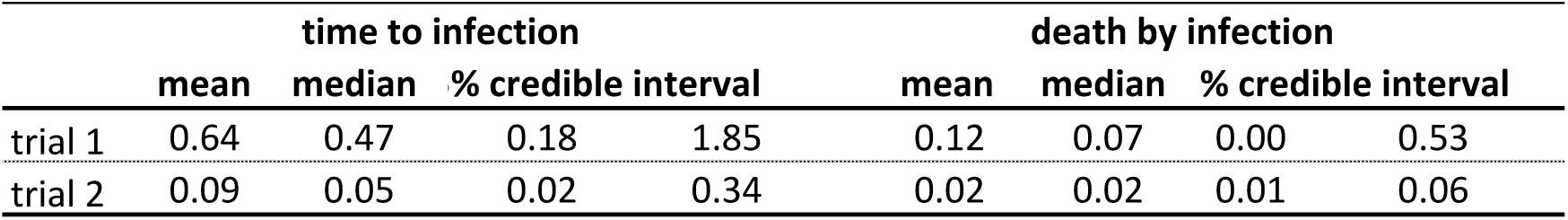
Estimates of variance components representing recipient family composition effects, included in the generalised linear mixed models for time to signs and time from signs to death

|  | time to infection |  |  |  | death by infection |  |  |  |
| --- | --- | --- | --- | --- | --- | --- | --- | --- |
|  | mean | median | % credible interval |  | mean | median | % credible interval |  |
| trial 1 | 0.64 | 0.47 | 0.18 | 1.85 | 0.12 | 0.07 | 0.00 | 0.53 |
| trial 2 | 0.09 | 0.05 | 0.02 | 0.34 | 0.02 | 0.02 | 0.01 | 0.06 |

**Figure 5.**
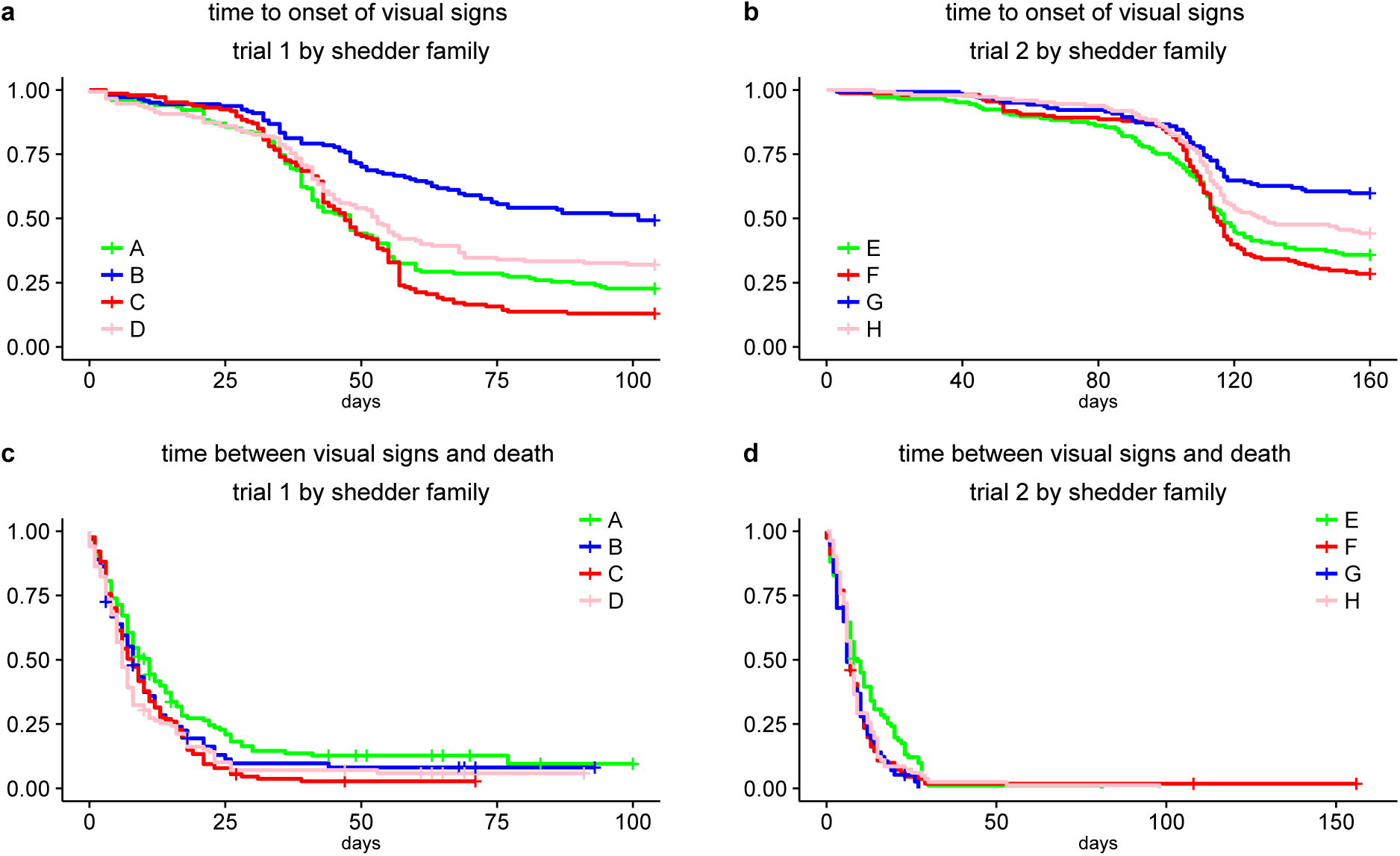
Effect of shedder fish family on recipient infection and survival post infection. Evolution of the epidemics and survival post infection in recipient fish pooled by families of shedder fish shared across recipients. **(a)** and **(b)** Kaplan-Meier curves for time to signs of recipient fish from trials 1 (T1) and 2 (T2) by shedder family. **(c)** and **(d)** Kaplan-Meier curves for time from signs to death for recipient fish by shedder family, also for both trials of the experiment. Number of recipient fish by shedder family for each of the graphs are presented in Supplementary table 4.

### Effect of recipient family composition on onset of disease and subsequent survival

To evaluate the effect of the genetic composition of the tank members (accounting for combined differences in recipient resistance, infectivity and tolerance) on disease and survival times, random effects representing recipient family composition were considered in all models (see Methods). Variance component estimates for recipient family composition are much larger in the models for time to onset of visual signs than in models for time between infection signs and death (Table 2), suggesting that the genetic composition of tank members played a strong role in disease spread but had little influence on survival post infection.

## Discussion

Host genetics has yet not been fully exploited to reduce the impact of infectious disease in animal populations. It is therefore crucial to understand and disentangle the effects of genetic diversity in different host defence mechanisms to disease. Here we presented a novel transmission experimental design which allows simultaneous genetic analyses of host resistance, tolerance and infectivity. Temporal epidemic data generated by applying our experimental design in an infectious disease model using family-structured fish allowed to dissect the different sources of genetic variation on disease prevalence and subsequent mortality. This contrasts with most genetic studies of infectious disease which focus on disease resistance alone, and often use binary mortality data at the end of the epidemic, thus not fully capturing the dynamic nature of infectious disease^41–43^.

Our study provided the first direct evidence that there are genetic differences in host infectivity and that the genetic make-up of a host can largely affect the survival of its group mates by affecting their risk of becoming diseased. The novel genetic differences found in host infectivity significantly explain variation in time to the appearance of visual infection signs, which in turn was found to be strongly related to survival. Host ability to transmit infections may also depend on the rate of contact between hosts or their excreted infectious material with susceptible individuals and on duration of the infection period^34,44^. The mode of the transmission of *Philasterides dicentrarchi* through contamination of water suggests constant rate of contact between susceptible fish and the parasites shed by hosts (see Methods). Moreover, we could not find significant family differences in shedder time to death in both trials (Supplementary Figure 2). This suggests that differences in infectivity are not simply explained by the fact that more infectious individuals shed longer nor that they are more tolerant to the disease. In addition, although higher infectivity can be also manifested by shedding higher quantity or more virulent infectious material^45^, there is no evidence for this in our study, as shedder family did not have a significant effect on recipient time from signs to death. Therefore, apart from the detected genetic variation in infectivity, we could not identify other significant epidemic factors that may explain variation in the ability to transmit the disease.

Our findings suggest that genetic variation in host resistance is larger than genetic variation in tolerance, and are in line with previous genetic studies of these traits for a variety of diseases and species^46–49^. However, the results may also partly depend on the trait definitions used in our study. Since time to infection can usually not be observed under natural transmission, we analysed resistance to onset of the disease, which may be different from resistance to becoming infected. Not developing disease after infection may be considered as an aspect of tolerance. Indeed, despite high sensitivity and specificity for onset of visual signs as a diagnostic test for infection by *P. dicentrarchi* (see Supplementary Table 1 and Methods), we found, through post mortem detection, parasites in 222 recipient fish (out of 1420 recipients that were post mortem examined – see Methods) which did not show signs of infection during the experiment. We surprisingly also detected neither parasites nor signs of infection in 53 out of the 180 artificially infected fish used in trial 2. These findings indicate that fish can express high levels of tolerance to infection that prevents development of infection signs, resulting in a minimum impact of the parasite on their health. This aspect of tolerance could however not be separated from resistance in our study.

Our experimental design tested in fish can be replicated in infection models with livestock species. The advantage of conducting these studies in aquaculture is the relative ease of creating many large families and generating large sample sizes. Conducting transmission experiments in fish or livestock model species may provide useful genetic information for studying complex disease traits such as resistance, tolerance and infectivity in the context of infectious diseases in human populations ^50^.

The family structure of the fish considered in our transmission experiment is expected to provide estimation of individual genetic risks for resistance, tolerance and infectivity^51^. In particular, under the assumption of infectivity being a heritable trait, it can be defined as an indirect genetic effect (IGE), also known as an associative or a social genetic effect^33,38,52,53^. Statistical models incorporating IGEs require populations structured into many isolated groups with related individuals in each of these groups^55,55^, which was the case in our experiment. We have recently extended IGE models to incorporate the dynamics of infection processes, and this extension can accurately estimate genetic risks as well as heritabilities and environmental effects for both infectivity and resistance^32^. However, further work is needed to adapt these models for infections that cause recovery or death (such as scuticociliatosis) and to incorporate genetic variation in tolerance to disease.

These IGE models have the advantage that they incorporate transmission between recipients into estimation of genetic risks for resistance, infectivity and tolerance, which was not explicitly considered in our study. In particular, these IGE models would provide infectivity genetic risk estimates for all individuals in a population, from which individuals with extreme risks can be identified as genetic superspreaders^32^. In the context of animal breeding, these genetic superspreaders have higher probability of generating offspring that would transmit most of the infections in a disease outbreak, and therefore preventing their occurrence would be an effective means to reduce disease prevalence in subsequent generations^56^.

In conclusion, our results imply that animals can potentially evolve three conceptually different types of response affecting their own and their group members’ chances of surviving infections. In particular, we demonstrate for the first time that there is significant genetic variation in in host infectivity. Our findings reveal new opportunities for devising more effective genetic disease control strategies by simultaneously exploiting genetic variation underlying host resistance, infectivity and tolerance to disease, rather than focusing only on disease resistance as measured by survival. As illustrated with a biological infection model in fish, genetic effects in resistance, tolerance and infectivity can be disentangled by a single experimental design. Broadening the scope of disease traits in genetic studies may open new avenues for identifying novel genes affecting disease transmission and survival as well as further understanding of mechanisms underlying infectious disease dynamics and evolution.

## Methods

### The scuticociliatosis transmission experiment in turbot

*Philasterides dicentrarchi* (*P. dicentrarchi*) is a histophagous ciliate causing scuticociliatosis in many fish species, in particular olive flounder and turbot. Scuticociliatosis is one of the most important parasitological problems in marine aquaculture worldwide^57^. In recent years, severe scuticociliatosis outbreaks with high mortality rates and devastating economic effects have been reported in East Asia (Korea, Japan, China), Europe (Spain, Portugal) and other regions of the world^37,58^. The parasite generates a severe systemic infection invading internal organs such as brain, gills, liver and intestine that generally results in the death of the host^59^.

Little is known about host genetic control of *P. dicentrarchi* infections. A QTL mapping study carried out using a controlled intracelomical challenge infection of turbot with a fixed high dose of virulent *P. dicentrarchi* strain, identified several genomic regions controlling dichotomous survival at the end of the experiment (28dpi) and survival time of infected fish, suggesting that survival is under genetic control^30^.

### Experimental facilities and turbot family structure

The farmed turbot used in the experiment were bred and reared in the experimental facilities of CETGA, Spain. The experiment complies with ethical regulations and was approved by the Regional Government of *Xunta de Galicia* (registered under the code ES150730055401/16/PROD.VET.047ROD.01). Due to space restrictions, the transmission experiments were carried out in two consecutive trials. Each trial comprised 900 fish (sex ratio 1:1) from 22 and 23 full-sibling families generated from 29 sires and 25 dams, for trials 1 and 2 respectively. The resulting full-sib families included eight and five paternal half-sibling families for respectively, trials 1 and 2 and seven maternal half-sib families in each of these trials. Parental fish were mostly unrelated, except for two paternal and three maternal half-sibs. Average body weights at the start of trials 1 and 2 were 32.1g (std. dev. 9.2g) and 91.6g (std. dev. 38.4g), respectively. Fish were tagged according to their full-sib family using elastomers, and fin clipping was used for individual identification.

### Distinction between shedder and recipient fish and their allocation across isolated experimental units

We assigned families in each trial into either shedder families (four per trial) or recipient families (18 per trial). Shedder families were inoculated with a virulent strain of *P. dicentrarchi* and then introduced into isolated tanks comprising non-infected recipient fish to seed the infection in the transmission experiment. Eventually, both infected shedders and recipients express infectivity, but artificial infection of shedders implies that shedder fish provide more accurate information about infectivity than recipient fish. To maximise genetic diversity, the eight shedder families were unrelated, i.e. shared no common sire or dam.

Each trial consisted of 36 independent transmission experiments (i.e. there is no between group transmission) carried out in 36 isolated 50-litre closed-circuit aerated tanks with constant water temperature of 17–18 °C and 0.36‰ salinity. To maximise statistical power for detecting and disentangle genetic variation in resistance and infectivity, we strategically distributed the shedder and recipient fish into the different tanks as follows (see also Fig. 1): each tank comprised 25 fish, with five fish from the same shedder family seeding the infection to 20 fish from 4 different recipient families (5 fish per recipient family), such that fish from each of the families were selected at random. Also, each shedder family seeded the infection in 9 different tanks (9 experimental replicates per shedder family, Fig. 1). To minimise confounding between shedder fish infectivity and genetic resistance and infectivity of recipient fish and therefore minimise the noise for detecting shedder family differences in infectivity, the same recipient family compositions were used across the 4 shedder families. This resulted in 9 different family compositions per trial, with each recipient family represented in 8 tanks and in 2 different family compositions. The sample size of 1800 fish and the family composition in tanks are in line with experimental design recommendations supported by theoretical results regarding studies of social genetic effects^60^ as well as simulation studies for detecting ideal experimental conditions for estimating infectivity genetic effects^32^.

### Infection of donors and transmission to recipients

Inoculation of shedder fish was carried out by intraperitoneal injection^59^ into the abdominal cavity of 200 µl of sterile physiological saline containing (50,000 and 56,000 ciliates were used in each of shedder fish from trials 1 and 2, respectively). After inoculation, infected shedder fish were immediately introduced into the tanks comprising initially only non-infected recipient fish and the epidemics were allowed to develop naturally. The trials were terminated at 104 and 160 days in trials 1 and 2, respectively. Fish still alive at the end of this period were killed with an overdose of anaesthetic. One of the tanks in trial 1 was discarded from the analysis due to a problem with oxygen supply. Also, due to technical difficulties in daily tracking of the disease status of all fish in the experiment, two fish from trial 1 and one fish from trial 2 had missing disease information and was also discarded from the analysis.

### Visual signs and histology

Visual infection signs included exophthalmia, colour change or depigmentation, visible lesions or abnormal swimming behaviour (no blinding method was used). Upon detection, dead fish were removed and both their brain and ascetic fluid collected from the body cavity were analysed to detect the presence of the scuticociliade. At the end of the experiment, all remaining fish were euthanized and screened for parasites.

### Statistical analyses

To compare infection and survival profiles associated with trials, tanks and families, we used Kaplan-Meier estimators for survival probability of developing disease and dying from disease using time to signs (with censoring given by surviving fish that did not show signs by the end of the experiment) and time to death, respectively. For fish that displayed visual infection signs, Kaplan-Meier estimators were also generated for time from signs to death. Fish that had signs but survived until the end of the experiment were censored in this case. The right-censoring mechanism considered was independent of the time to signs and time to death observed in recipient fish, a crucial assumption for calculation of Kaplan-Meier curves and log-rank tests.^61^

Daily counts of recipient fish with visual infection signs in each tank were used to estimate the effect of shedder fish family on time to infection. For a tank i and day t_j_ of the experiment, the number of fish with visual signs, represented by C_i_(t_j_) was assumed to follow a binomial distribution with parameters given by the number of susceptible individuals at day t_j_ in tank i, represented by S_i_(t_j_), and p_j_, which is the probability of a fish showing visual signs at t_j_ given it was susceptible prior to that day. This conditional probability is, by definition, the disease hazard at day t_j_ ^39,40^ and can be modelled using a generalised linear mixed model with a complementary log-log link function given by

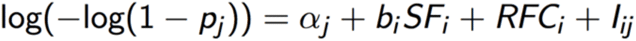

where α_j_ is the day-specific intercept representing a baseline hazard, SF_i_ is the shedder family at tank i with effect b_i_ and I_ij_ is an offset representing the proportion of infected fish at tank i and day t_j_. Individual family effects were not incorporated in this tank-day level model as each tank comprised four recipient families. Therefore, RFC_i_ is a random effect representing recipient family composition at tank i such that its covariance matrix incorporates family relationships within and between the compositions^62^. A similar model was considered to estimate the shedder family effect on time to death by infection by assuming that the daily number of recipient fish that died to infection at tank i, represented by R_i_(t_j_), follows a binomial distribution with parameters given by the number of diseased fish alive at day t_j_ in tank i (S_i_(t_j_)) and π_j_, which is the probability of a fish dying by infection t_j_ given it was alive and displayed signs prior to that day. Both models were also fitted ignoring shedder family effect and compared to the full models using the Watanabe-Akaike information criterion^63^. Bayesian inference for model parameters was based on the Hamiltonian Monte Carlo algorithm to generate samples of posterior distributions, implemented in the Stan package of the R statistical software^64^.

## Supporting information

## Code availability

The computer code used in this study are available from the corresponding author upon reasonable request.

## Data Availability

The data that support the findings of this study are available from the *Centro Tecnologico del Cluster de la Acuicultura de Galicia* (CETGA), but restrictions apply to the availability of these data, which were used under license for the current study, and so are not publicly available. Data are however available from the corresponding author upon reasonable request and with permission of CETGA.

## Acknowledgements

This work was carried out with funding from the Biotechnology and Biological Sciences Research Council Institute Strategic Programme grant ISP1 (to O.A., J.A.W., R.H and A.B.D.-W.) and the European Union’s Seventh Framework Programme (KBBE.2013.1.2-10) under grant agreement n° 613611 (all authors).

## Author contributions

All the authors conceived the experimental design. S.C. carried out the experiment. O.A. carried out the statistical analyses. O.A and A.B.D.-W wrote the first draft of the manuscript and all the authors contributed to discussing the results and editing the manuscript.

