## Supplementary material for "Genetic differences in host infectivity affect disease spread and survival in epidemics"

Osvaldo Anacleto[1], Santiago Cabaleiro[2], Beatriz Villanueva[3], María Saura[3], Ross D. Houston[1], John A. Woolliams[1], Andrea B. Doeschl-Wilson[1]
1The Roslin Institute and Royal (Dick) School of Veterinary Studies, University of Edinburgh, UK
[2]Centro Tecnológico del Cluster de la Acuicultura (CETGA), Spain
[3]Departamento de Mejora Genética Animal, INIA, Spain

**Turbot fish infected by *Philasterides dicentrarchi* can be accurately identified through visual signs.** Shedder fish successfully transmitted the parasite to recipients. In our experiment, 766 out of 1417 recipient fish showed signs of infection and *Philasterides dicentrarchi* was detected by post-mortem inspection in 94% of these fish (Figure 2). Although the parasite was detected post-mortem in 34% of recipients with no visual signs of infection (see Discussion), high true positive proportions (sensitivity) and true negative proportions (specificity) when considering these signs as a diagnostic test suggest that they are reliable indicators of presence of Philasterides dicentrarchi in fish (Supplementary Table 1, also see Methods). Infection by *Philasterides dicentrarchi* was the only cause of natural death during the trials, and 945 out of 1417 recipient fish died during the experiment.

**Supplementary Table 1:** Sensitivity and specificity of onset of visual signs to detect presence of *Philasterides dicentrarchi* in Turbot Fish

| | **sensitivity (true positive)** | **specificity (true negative)** |
|---|---|---|
| shedders (n=355) | 0.81 (239/294) | 0.89 (54/61) |
| recipients (n=1417) | 0.77 (723/945) | 0.91 (429/472) |

Post-mortem detection of *Philasterides dicentrarchi* was used as gold standard to verify the quality of the onset of visual signs as infection diagnostic test.

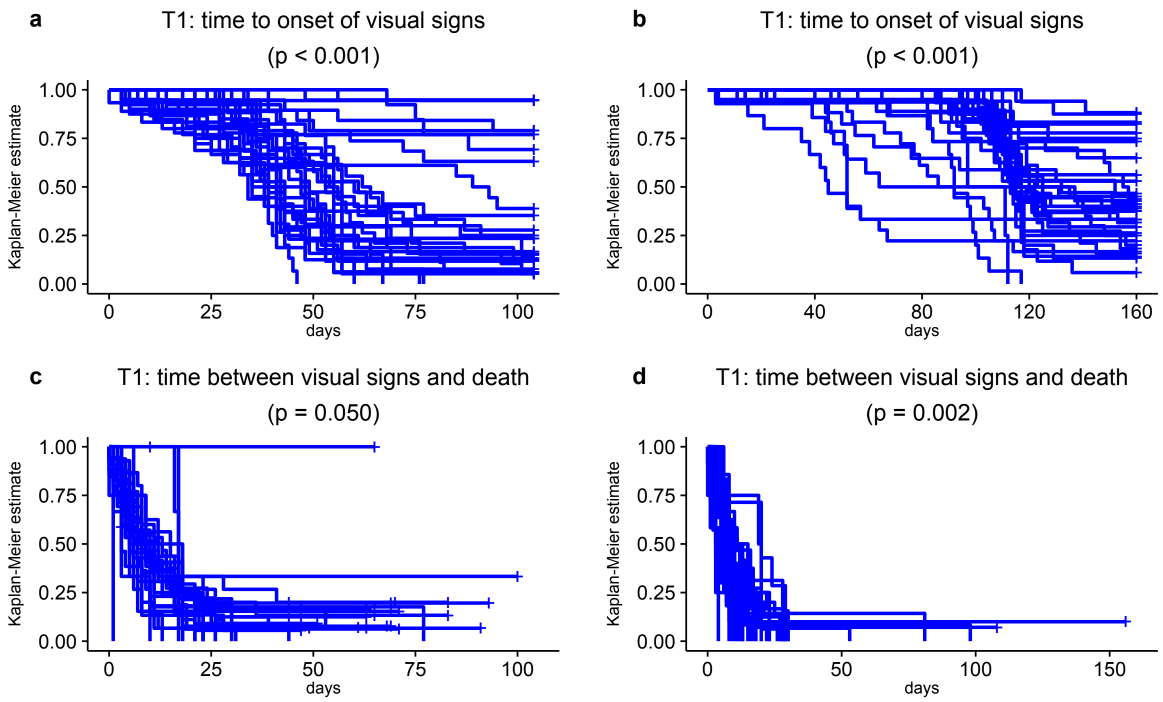


**Supplementary Figure 1. Tank differences in onset of visual signs and survival post disease.** Evolution of disease caused by *Philasterides dicentrarchi* (a-b) and survival post disease (c-d) in all families of recipient fish in trials 1 (T1) and 2 (T2) of the transmission experiment. The curves were obtained through tank-based Kaplan-Meier plots for time to signs (a-b) and time from signs to death (c-d). P-values were calculated using the two tailed log-rank test for detecting family differences in Kaplan-Meier estimates.

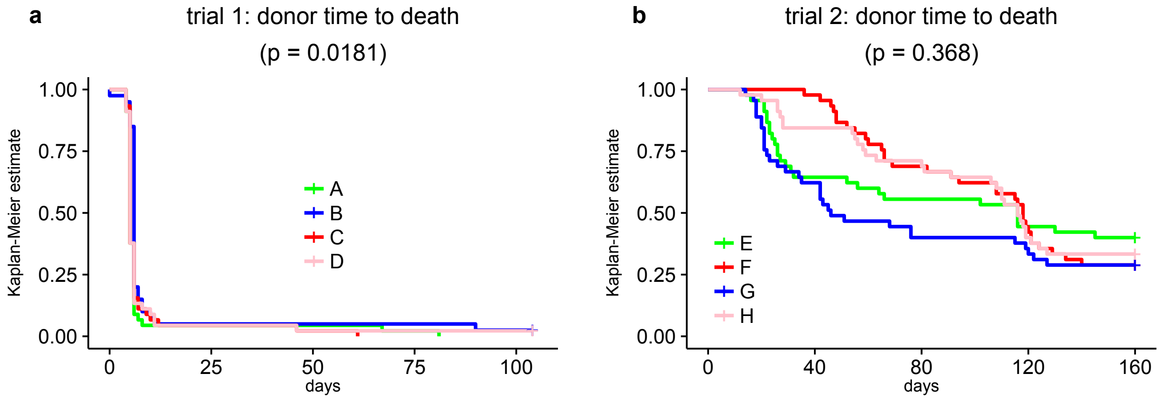


**Supplementary Figure 2. Survival of shedder fish families.** Kaplan-Meier curves for time to death of shedder fish for trials 1 (a) and 2 (b), by family. Most infective shedder families were C and F for trials 1 and 2, respectively. Least infective shedder families were B and G for trials 1 and 2, respectively. P-values were calculated using the two tailed log-rank test for detecting family differences in Kaplan-Meier estimates

**Supplementary Table 2.** Number of fish from each of the trial 1 recipient families shown in Kaplan-Meier plots in Figures 4a and 4c.

| family | time to disease signs | | | time from signs to death | | |
|---|---|---|---|---|---|---|
| | **event** | **censored** | **total** | **event** | **censored** | **total** |
| 7 | 12 | 27 | 39 | 12 | 0 | 12 |
| 8 | 30 | 3 | 33 | 26 | 4 | 30 |
| 10 | 28 | 8 | 36 | 26 | 2 | 28 |
| 11 | 26 | 9 | 35 | 23 | 3 | 26 |
| 12 | 26 | 10 | 36 | 24 | 2 | 26 |
| 13 | 28 | 6 | 34 | 27 | 1 | 28 |
| 14 | 21 | 4 | 25 | 18 | 3 | 21 |
| 15 | 25 | 9 | 34 | 21 | 4 | 25 |
| 16 | 23 | 11 | 34 | 23 | 0 | 23 |
| 18 | 25 | 2 | 27 | 24 | 1 | 25 |
| 19 | 21 | 15 | 36 | 18 | 3 | 21 |
| 20 | 13 | 14 | 27 | 12 | 1 | 13 |
| 22 | 26 | 7 | 33 | 26 | 0 | 26 |
| 28 | 26 | 3 | 29 | 26 | 0 | 26 |
| 30 | 13 | 20 | 33 | 11 | 2 | 13 |
| 32 | 25 | 7 | 32 | 21 | 4 | 25 |
| 33 | 27 | 8 | 35 | 24 | 3 | 27 |
| 39 | 25 | 9 | 34 | 20 | 5 | 25 |
| **Total** | **420** | **172** | **592** | **382** | **38** | **420** |

**Supplementary Table 3.** Number of fish from each of the trial 2 recipient families shown in Kaplan-Meier plots in Figures 4b and 4d.

| family | time to disease signs | | | time from signs to death | | |
|---|---|---|---|---|---|---|
| | event | censored | total | event | censored | total |
| 9 | 16 | 14 | 30 | 16 | 0 | 16 |
| 36 | 21 | 11 | 32 | 21 | 0 | 21 |
| 38 | 14 | 18 | 32 | 14 | 0 | 14 |
| 42 | 17 | 9 | 26 | 17 | 0 | 17 |
| 43 | 19 | 18 | 37 | 19 | 0 | 19 |
| 45 | 18 | 13 | 31 | 17 | 1 | 18 |
| 47 | 29 | 5 | 34 | 28 | 1 | 29 |
| 48 | 25 | 9 | 34 | 25 | 0 | 25 |
| 52 | 16 | 20 | 36 | 16 | 0 | 16 |
| 53 | 19 | 10 | 29 | 18 | 1 | 19 |
| 54 | 16 | 21 | 37 | 15 | 1 | 16 |
| 55 | 19 | 11 | 30 | 19 | 0 | 19 |
| 56 | 23 | 13 | 36 | 23 | 0 | 23 |
| 58 | 17 | 18 | 35 | 16 | 1 | 17 |
| 59 | 19 | 10 | 29 | 19 | 0 | 19 |
| 60 | 19 | 15 | 34 | 19 | 0 | 19 |
| 63 | 9 | 23 | 32 | 9 | 0 | 9 |
| 70 | 26 | 8 | 34 | 26 | 0 | 26 |
| **Total** | **342** | **246** | **588** | **337** | **5** | **342** |

**Supplementary Table 4.** Number of recipient fish exposed to each of the eight shedder families shown in Kaplan-Meier plots in Figure 5.

| shedder family | time to disease signs | | | time from signs to death | | |
|---|---|---|---|---|---|---|
| | event | censored | total | event | censored | total |
| 23 | 118 | 35 | 153 | 102 | 17 | 119 |
| 24 | 73 | 71 | 144 | 63 | 10 | 73 |
| 25 | 127 | 18 | 145 | 123 | 4 | 127 |
| 27 | 101 | 48 | 149 | 94 | 7 | 101 |
| 31 | 93 | 51 | 144 | 92 | 1 | 93 |
| 44 | 111 | 45 | 156 | 107 | 3 | 110 |
| 46 | 57 | 85 | 142 | 56 | 1 | 57 |
| 51 | 82 | 65 | 147 | 82 | 0 | 82 |
| **Total** | **762** | **418** | **1180** | **719** | **43** | **762** |
